# Bioinformatic analysis of shared B and T cell epitopes amongst relevant coronaviruses to human health: Is there cross-protection?

**DOI:** 10.1101/2020.07.14.202887

**Authors:** Diana Laura Pacheco-Olvera, Stephanie Saint Remy-Hernández, Ernesto Acevedo-Ochoa, Lourdes Arriaga-Pizano, Arturo Cérbulo-Vázquez, Eduardo Ferat-Osorio, Tania Rivera-Hernández, Constantino López-Macías

## Abstract

Within the last 30 years 3 coronaviruses, SARS-CoV, MERS-CoV and SARS-CoV-2, have evolved and adapted to cause disease and spread amongst the human population. From the three, SARS-CoV-2 has spread world-wide and to July 2020 it has been responsible for more than 11 million confirmed cases and over half a million deaths. In the absence of an effective treatment or vaccine, social distancing has been the most effective measure to control the pandemic. However it has become evident that as the virus spreads the only tool that will allow us to fully control it is an effective vaccine. There are currently more than 150 vaccine candidates in different stages of development using a variety of viral antigens, with the S protein being the most targeted antigen. Although some new experimental evidence suggests cross-reacting responses between coronaviruses are present in the population, it remains unknown whether potential shared antigens between different coronaviruses could provide cross-protection. Given that coronaviruses are emerging pathogens and continue to represent a threat to global health, the development of a SARS-Cov-2 vaccine that could provide ‘universal’ protection against other coronaviruses should be pushed forward. Here we present a thorough review of reported B and T cell epitopes shared between SARS-CoV-2 and other relevant coronaviruses, in addition we used web-based tools to predict novel B and T cell epitopes that have not been reported before. Analysis of experimental evidence that is constantly emerging complemented with the findings of this study allow us support the hypothesis that cross-reactive responses, particularly those coming from T cells, might play a key role in controlling infection by SARS-CoV-2. We hope that with the evidence presented in this manuscript we provide arguments to encourage the study of cross-reactive responses in order to elucidate their role in immunity to SARS-CoV-2. Finally we expect our findings will aid targeted analysis of antigen-specific immune responses and guide future vaccine design aiming to develop a cross reactive effective vaccine against respiratory diseases caused by coronaviruses.

## 1 Introduction

At the end of December 2019, the Chinese government emitted a sanitary alert regarding a novel coronavirus which was responsible of a significant number of severe respiratory infections. SARS-CoV-2 was identified as the etiological agent causing what was later designated as coronavirus disease (COVID-19). Until July 2020, SARS-CoV-2 has caused more than 11 million confirmed infections and over half a million deaths world-wide (1). In the previous 30 years we have witnessed the emergence of three coronaviruses that crossed the species barrier and gained the availability to infect the human host. SARS-CoV, MERS-CoV and now SARS-CoV-2 have evolved from their original animal host and adapted to cause severe respiratory disease in humans (2). Genetic studies have shown that the newly emerged SARS-CoV-2 shares 96% genetic similarity with a bat betacoronavirus belonging to the *Sarbecovirus* subgenus, while it shares 79% and 50% genetic similarity with SARS-CoV and MERS-CoV, respectively (3). Closer and increasing human interaction with wildlife is key in the evolution of these coronaviruses, thus it is expected that more outbreaks of emergent viruses as such will continue to occur in the future, representing a constant threat to human health (4). On the other hand, other human coronaviruses, HCoV-HKU1 (HKU), HCoV-NL63 (NL63), HCoV-OC43 (OC43) and HCoV-229E (229E), circulate amongst the human population in a seasonal manner causing non-severe respiratory disease (2). Immune responses against infections with outbreak or seasonal coronaviruses differ significantly. Evidence suggests that immunity to seasonal coronaviruses is short-lasting with antibodies quickly decreasing after infection (5), and controlled infection of human volunteers suggests infection does not generate protective immunity against re-infection (6). On the other hand, it has been reported that patients who recovered from infection with SARS-CoV carry circulating antibodies up to 12 years after the infection occurred, although the neutralising activity of such antibodies was not investigated (7).

As the current pandemic unfolds, it has become evident that the only tool that will allow humanity to control the virus spread is the development of a safe and effective vaccine. To the date there are more than 150 vaccine candidates in different phases of preclinical and clinical development (https://www.covid-19vaccinetracker.org/) with most candidates targeting the spike glycoprotein that mediates viral entry to the host cell via binding to angiotensin-converting enzyme 2 (ACE2) (8). In order to develop a successful vaccine it is imperative to understand the antigenic nature of the main target, the spike glycoprotein. Moreover, limited information is available regarding the antigenic similarity of SARS-CoV-2 glycoprotein and other coronaviruses whether causing seasonal disease or outbreaks. Historically neutralising antibodies have been regarded as a correlate for protection following vaccination or infection, and even though there are some studies in the literature, little attention has been given to study T cell responses and their role in protection (9, 10). Here we present a compilation of potential B and T cell epitopes found in the SARS-CoV-2 spike glycoprotein and discuss their potential role in generating cross-reactive immune responses against other human coronaviruses. We aim that this study will support and strengthen the hypothesis that T cell responses play a key role in immunity to SARS-CoV-2 including pre-existing responses to other seasonal coronaviruses. In addition we hope that the epitope compilation will aid the rational design of a vaccine that could provide cross-protection against other coronaviruses that currently circulate amongst the human population or even other coronaviruses that might emerge in the future from other animal species with pandemic potential.

## 2 Materials and Methods

### 2.1 Epitope prediction

Genome sequences for human coronaviruses HCoV-HKU1 (HKU), HCoV-NL63 (NL63), HCoV-OC43 (OC43), HCoV-229E (229E), MERS-CoV, SARS-CoV and SARS-CoV-2 were accessed via NCBI, Genbank accession numbers YP_173238.1, YP_003767.1, YP_009555241.1, ABB90529.1, AKN11075.1, AAU04646.1 and QHR63290.2 respectively. Protein multiple sequence alignments were performed using Clustal Omega (11). Prediction of linear B cell epitopes was performed via the Immune Epitope Database (IEDB) website (https://www.iedb.org/) using the Bepipred Linear Epitope Prediction algorithms V1.0 and V2.0 (12) Threshold values of 0.35 (corresponding specificity >0.49 and sensitivity <0.75) and 0.55 (corresponding specificity > 0.817 and sensitivity <0.292) were used with versions 1.0 and 2.0, respectively. Epitopes were chosen based on their IC50 binding values of <50 (high affinity) and <500 (medium affinity). In order to determine if the predicted B cell epitopes were exposed in the protein surface, the surface accessibility and secondary structure, NetSurfP-2.0 was used (13). Prediction of conformational B cell epitopes was performed using the ElliPro and DiscoTope 2.0 tools from IEDB website (14, 15). Discontinuous B-cell epitopes were predicted via the DiscoTope 2.0 server tool in IEDB with a default threshold of −3.7 (corresponding specificity > 0.75 and sensitivity < 0.47), based on the 3-dimensional (3D) structures of the SARS-CoV-2 S protein (PDB ID: 6VYB, A chain). For T cell epitope prediction with MHC-I restriction, the MHC-I Binding Predictions (NetMHCpan EL 4.0 method) (16) and Class I Immunogenicity (17) tools from the IEDB website were used. T cell epitope selection was done based on IC50 affinity scores (<500) and immunogenicity scores (>-1 and <1). For T cell epitope prediction with MHC-II restriction, we used the IEDB-recommended algorithm 2.22. The prediction was carried out for 3 HLA-DR alleles (HLA-DRB1*01:01, HLA-DRB1*04:01, HLA-DRB1 *07:01), 8 HLA-DP alleles (HLA-DPA1*01:03/DPB1*03:01, HLA-DPA1*02:01/DPB1*02:01, HLA-DPA1*02:02/DPB1*02:02, HLA-DPA1*03:01/DPB1*23:01) and 6 HLA-DQ alleles (HLA-DQA1*05:01/DQB1*03:01, HLA-DQA1*01:01/DQB1*05:01, HLA-DQA1*03:01/DQB1*03:02) with a limit of 15 amino acid peptide length and a median consensus percentile of prediction threshold ≤20. T cell epitope selection was done based on IC50 affinity scores (<500). Epitope identity between SARS-CoV-2and other coronaviruses was calculated using the EMBOSS Needle pairwise sequence alignment tool (11).

### 2.2 Structural Modelling

To provide a graphical representation of the epitopes, we used the structural model of the full-Length SARS-CoV-2 spike glycoprotein (ID: 6VSB_1_1_1) (18, 19). The 3D structures were built and analysed using PyMOL^®^ software (Schrödinger LLC. Molecular Graphics System (PyMoL) Version 1.80 LLC, New York, NY. 2015). The basic local alignment search tool online (https://blast.ncbi.nlm.nih.gov/Blast.cgi) was used to assess the position of the predicted peptides in the glycoprotein sequence.

## 3 Results

Using web-based tools we carried out a bioinformatic analysis of the SARS-CoV-2 glycoprotein sequence, in order to identify novel B and T cell epitopes with potential to be used as vaccine targets. In order to do so we followed the strategy shown in figure 1, where we show the tools and parameters chosen to carry out the analysis.

**Figure 1.**
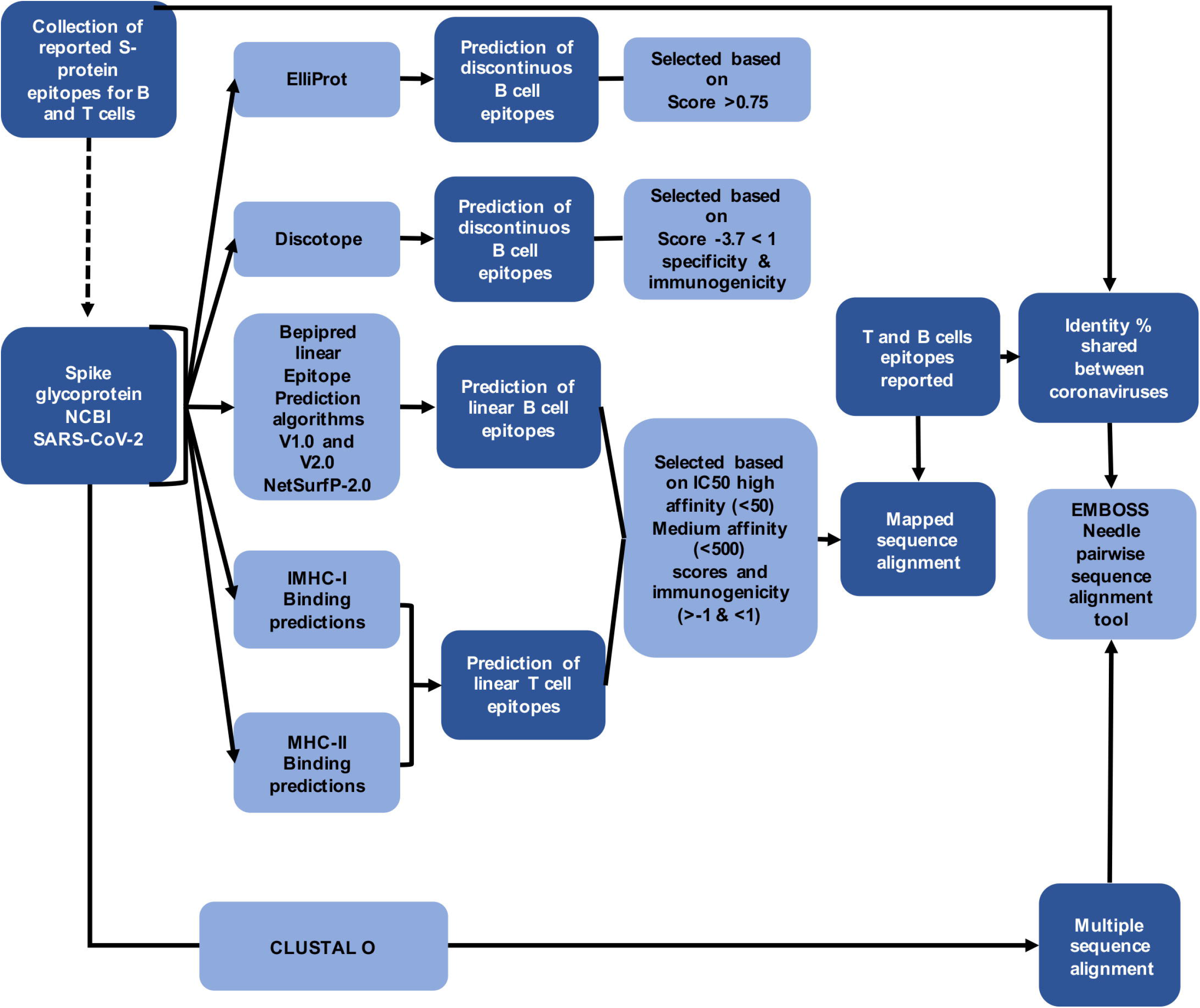
Workflow chart of the strategy followed in this study.

### 3.1 B cell epitope prediction

We carried out a literature review of studies reporting epitope mapping of the spike glycoprotein form SARS-CoV-2 and were able to compile a total of 41 potential B cell epitopes (Supplementary Table 1) that were identified through bioinformatic analysis by other research groups (20–23). By using BepiPred 1.0 and 2.0 we predicted 101 epitopes from which 41 were selected based on their score and surface accessibility. In total, we compiled 82 potential B cell epitopes (from our study and previous reports) from which 14 were both predicted in this study and reported by other groups and 27 were novel epitopes predicted in this study. Epitopes that coincided with previously reported epitopes or had overlapping residues, were merged into one peptide which covers all reported and predicted epitopes and are shown in bold letters in Supplementary Table 1. To visualise the position of the compiled epitopes we used an optimised 3D model of the SARS-CoV-2 spike glycoprotein as a monomer (Supplementary figure 1) and in its trimeric conformation (Figure 2) (19). B cell epitopes reported in the literature are distributed in all domains of the spike glycoprotein similarly to epitopes predicted in this study (Supplementary figure 1A & B). The trimeric model shows that linear B cell epitopes predicted in this study are highly abundant in the top region the trimer compared to reported epitopes (Figure 2A & B). These regions comprise the S1 polypeptide which contains the N-terminal domain (NTD), the receptor binding domain (RBD) and the receptor binding motif (RBM). A multiple sequence alignment of the SARS-CoV-2 spike glycoprotein against the glycoprotein sequences from other human coronaviruses (SARS-CoV, MERS-CoV, HKU1, NL63, OC43 and 229E) allowed us to identify conserved epitopes amongst these coronaviruses that could be target of cross-reacting antibodies. We were able to identify 7 different epitopes that share certain identity between the seven coronaviruses (Table 1). Most of the conserved epitopes are situated on the heptad repeat 1 (HR1) (Supplementary figure 1C) with some of them facing the inner part of the trimeric structure (Figure 2C). Table 1 shows the sequences of the conserved peptides with the percentage of identity between the SARS-CoV-2 sequence and the other six coronaviruses. Only one of the seven peptides (VEAEVQIDRLITGRLQSL) showed a percentage of identity higher than 60% between SARS-CoV-2 and all the other coronaviruses sequences. As expected, epitopes between SARS-CoV and SARS-CoV-2 shared the highest identity percentages between coronaviruses.

**Figure 2.**
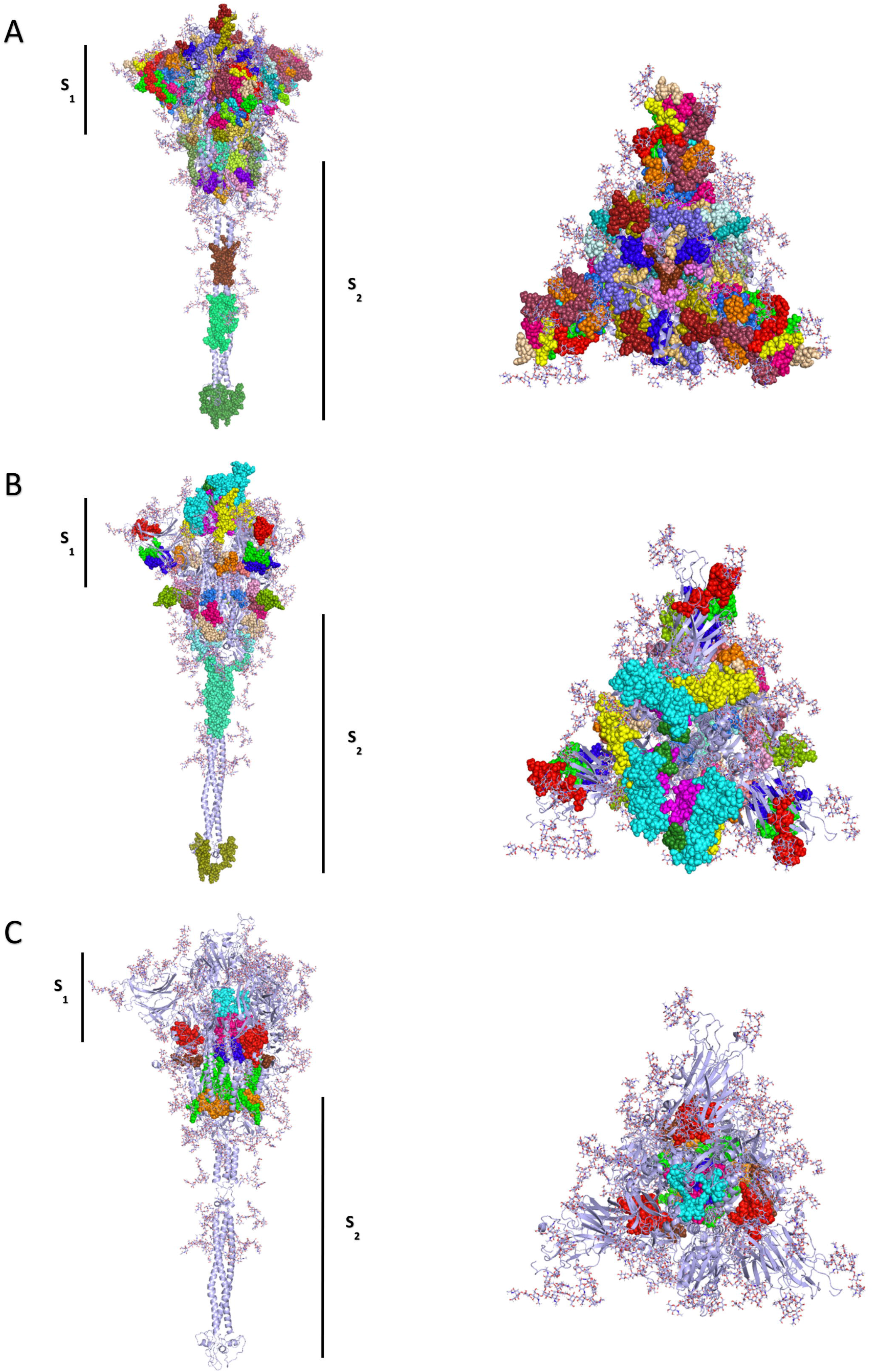
Linear B cell epitopes of the SARS-CoV-2 spike glycoprotein displayed in the trimer representation, side and top view. Epitopes reported in the literature (A), epitopes predicted in this study with IEDB tools (B) and epitopes that are shared with other coronaviruses predicted or reported (C).

**Table 1.**
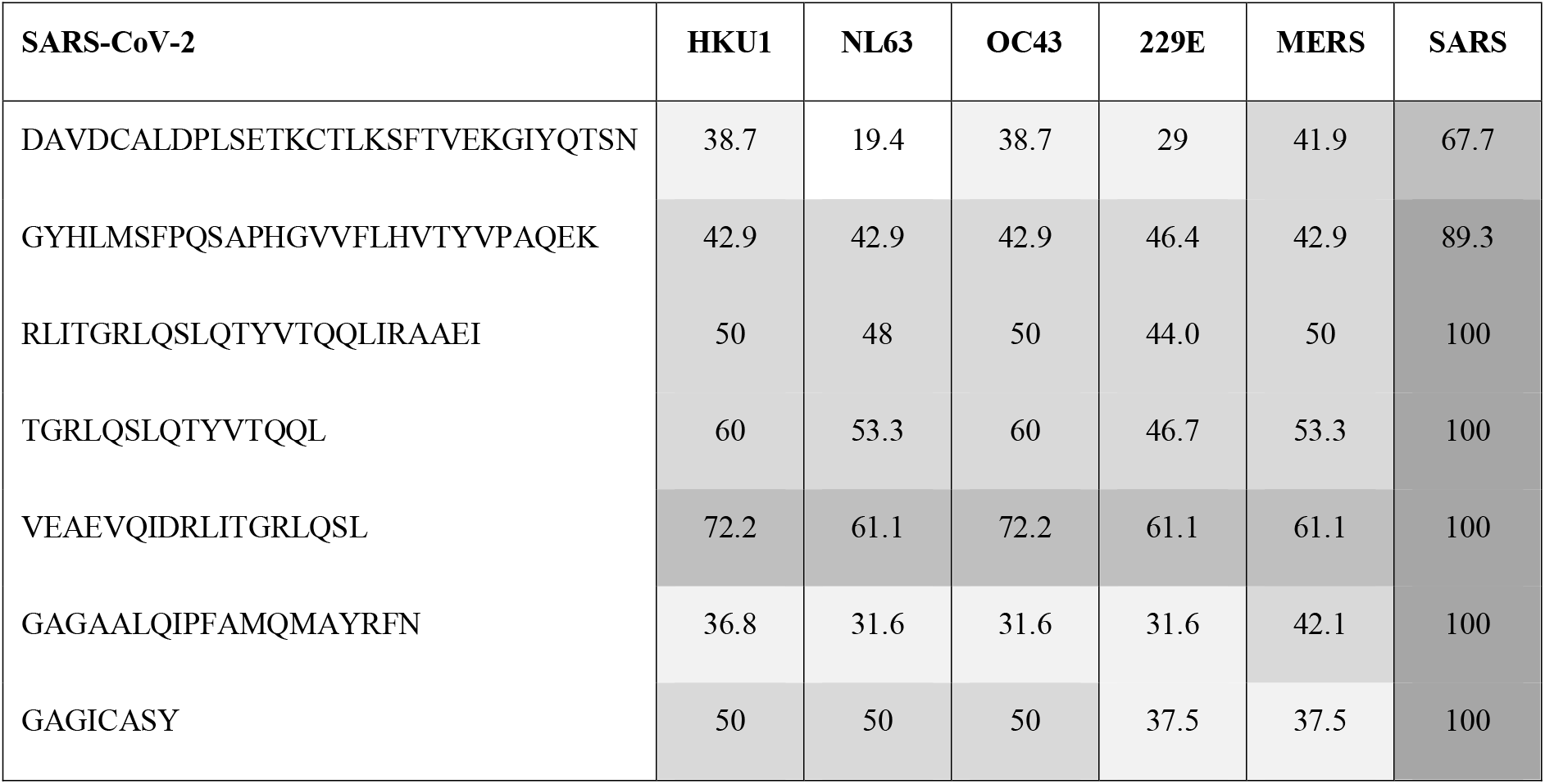
SARS-CoV-2 reported linear B cell epitopes identity (%) shared with other coronaviruses.

Using the ElliPro and DiscoTope tools from the IEDB website we predicted potential discontinuous epitopes on the SARS-CoV-2 spike glycoprotein. Prediction using ElliPro yielded 12 discontinuous epitopes (Figure 3A) of which 3 have residues belonging to the RBD and residues belonging to the NTD. A single epitope contains residues in the region of the fusion peptides (FP) and in HR1 (Figure 3A). The remaining 8 epitopes have residues situated in the transitional region between the S1 and S2 subunits (Figure 3A). Three epitopes predicted by ElliPro contained residues from 2 different monomers, these epitopes are shown in colours red, blue and green in figure 3A. In addition, one epitope shown in yellow in figure 3A, contained residues from each of the 3 monomers from the spike glycoprotein trimer, suggesting the presence of a highly antigenic region within the trimeric structure. Prediction of conformational epitopes using DiscoTope 2.0 resulted in 967 surface residues from which 55 were chosen based on a sensitivity of 0.47 and specificity of 0.75. The selected residues are distributed mainly in 4 domains (Figure 3B). In particular, one residue (N282) belongs to the NTD, and the RBD contains 28 of the predicted residues with 27 of these situated in the RBM (Figure 3B). Within the S2 region, three residues (P793, I794 and P809) are located in the fusion peptide (FP) and another three residues (N914, Y917 and E918) fall within the HR1. The predicted residues are shown in Figure 3B, where predicted residues with a high specificity (91 to 100%) are shown in red, the residues with medium specificity (86 to 90%) are shown in yellow and residues with a low specificity (75 to 85%) are shown in blue.

**Figure 3.**
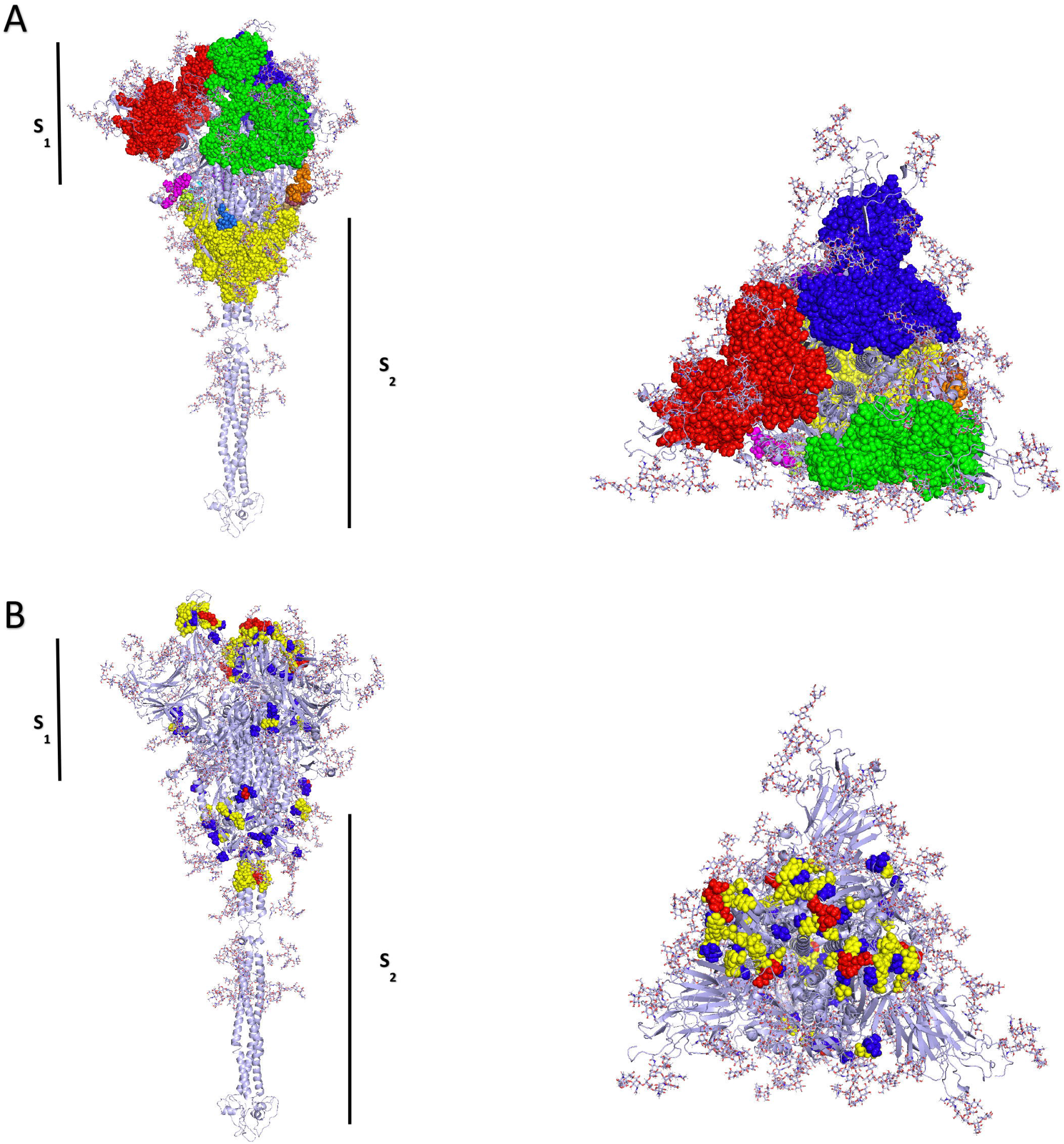
Conformational B cell epitopes of the SARS-CoV-2 spike glycoprotein obtained using in the (A) ElliPro and (B) DiscoTope prediction tools, side and top view. (A) The ElliPro prediction shows the major epitopes between chain A and chain B are shown in red, for chain B and C are shown in green and finally for chain A and C are shown in yellow. (B) The DiscoTope prediction the residues with a high specificity (91 to 100%) are shown in red, the residues with medium specificity (86 to 90%) are shown in yellow and residues with a low specificity (75 to 85%) are shown in blue.

### 3.2 T cell epitope prediction

From reports in the literature we compiled 107 peptide sequences reported to be CD8 and CD4 T cell epitopes (Figure 4A and Supplementary figure 2A) (21, 23, 24). For prediction of peptides with MHC-I restriction we used NetMHCpan EL 4.0 method, while peptides with MHC-II restriction the IEDB-recommended algorithm 2.22 was used. HLA selection was done following methodologies previously reported (25) (26) and used for prediction with the TepiTool resource (27). In addition, 4 additional alleles (1 for HLA-I and 3 for HLA-II) were used for prediction given that they are present at a high frequency in the Mexican population and also have a significant frequency worldwide. Additional alleles were obtained from the allelic frequency database (http://www.allelefrequencies.net/default.asp), where the search criteria was the North American region, all groups from the Mexican population within capital cities across the country. Using these parameters we were able to predict a total of 66 possible T cell epitopes (Figure 4B and Supplementary figure 2B), from which 30 have predicted binding to MHC-I, 32 have predicted binding to MHC-II and 4 are predicted to be promiscuous T cell epitopes (Supplementary Table 2). Reported epitopes found in the literature can be found all throughout the spike glycoprotein structure (Figure 4A). In contrast, predicted epitopes are present throughout the structure with the exception of a noticeable gap in the transmembrane domain (TD) (Figure 4B). Through a multiple sequence alignment we identified 38 epitopes (Figure 4C and Supplementary figure 2C) that shared certain % of identity between the SARS-CoV-2 spike glycoprotein and the other human coronaviruses (SARS-CoV, MERS-CoV, HKU1, NL63, OC43 and 229E). Table 2 shows the 38 conserved epitopes with the percentage of identity between the SARS-CoV-2 sequence and the other six coronaviruses. Several epitopes shared more than 60% identity between SARS-CoV-2 and all the other human coronaviruses, being epitopes from SARS-CoV the ones with the highest percentages of identity with SARS-CoV-2.

**Figure 4.**
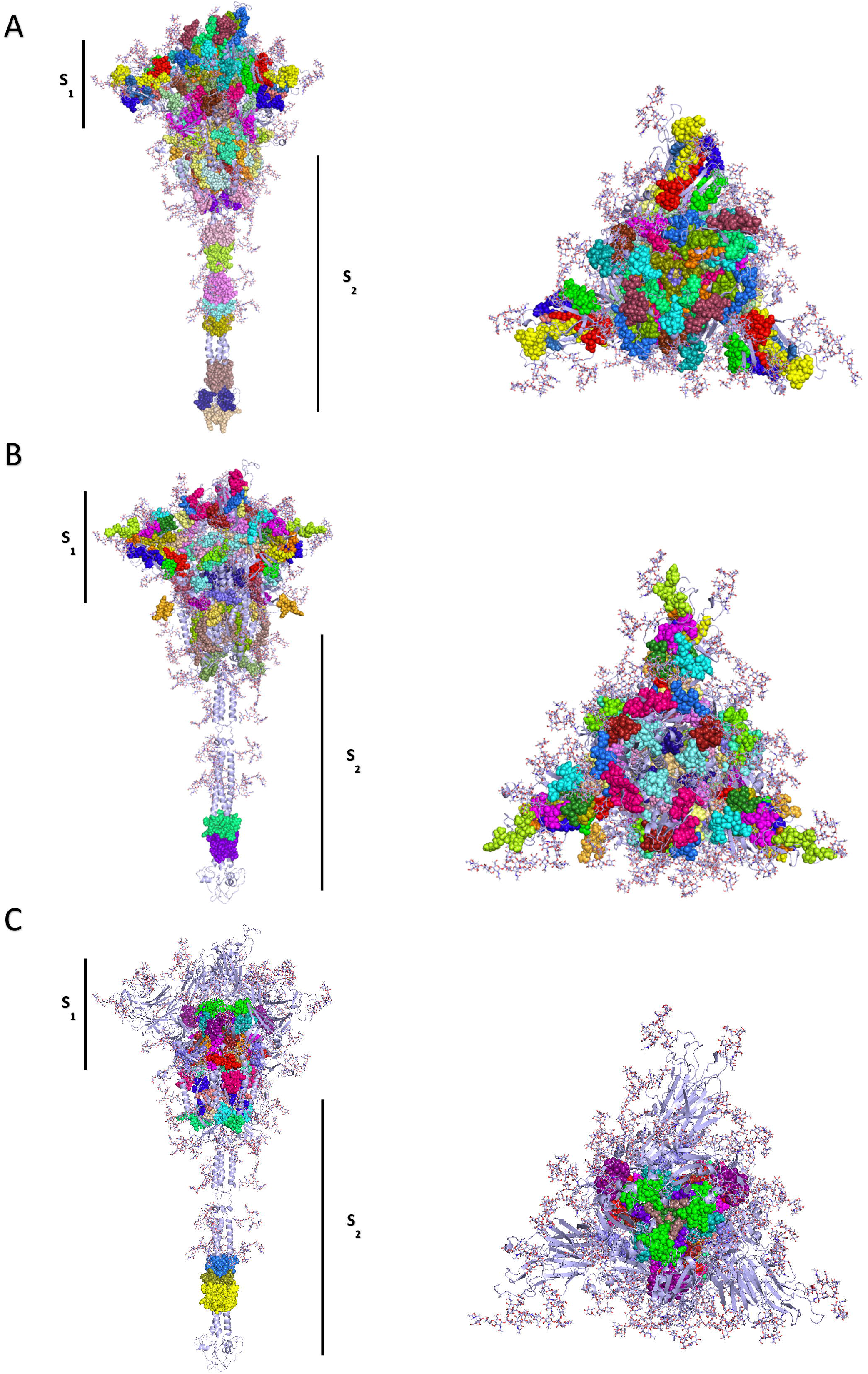
Epitopes of the SARS-CoV-2 spike glycoprotein restricted to MHC-I and MHC-II displayed in the trimer representation, side and top view. Epitopes reported in the literature (A), epitopes predicted in this study using IEDB tools (B) and epitopes that are shared with other coronaviruses predicted or reported (C).

**Table 2.**
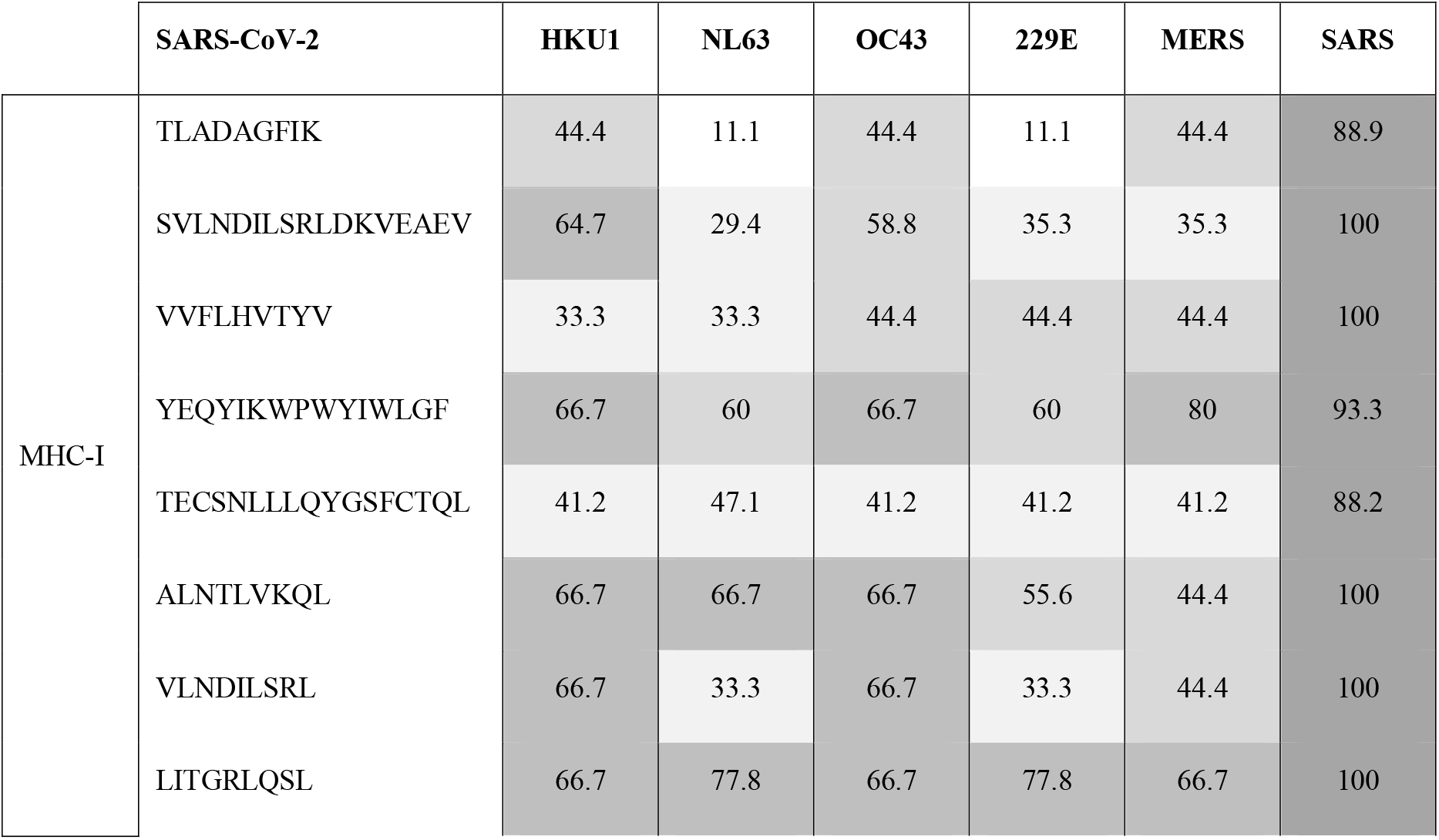
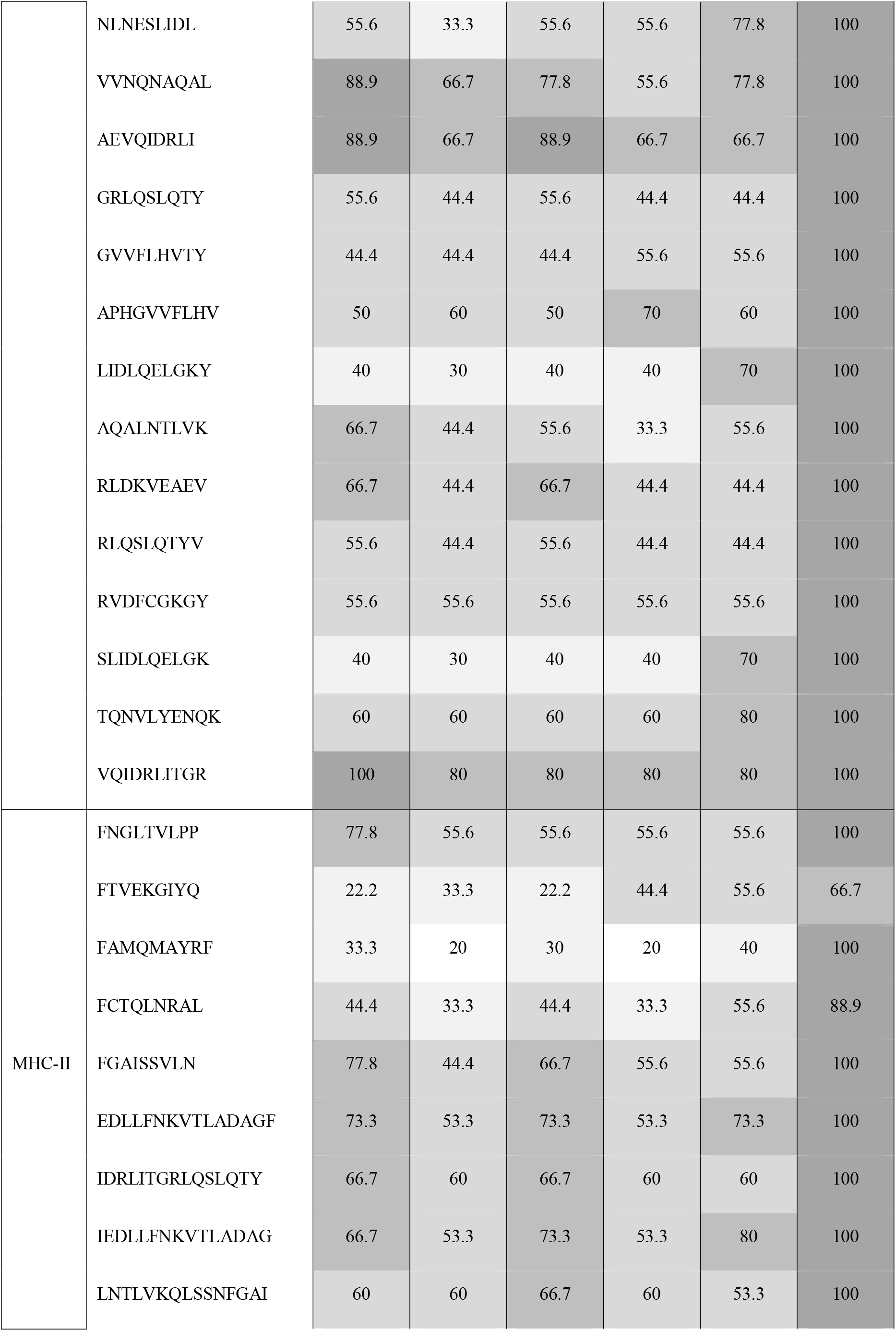
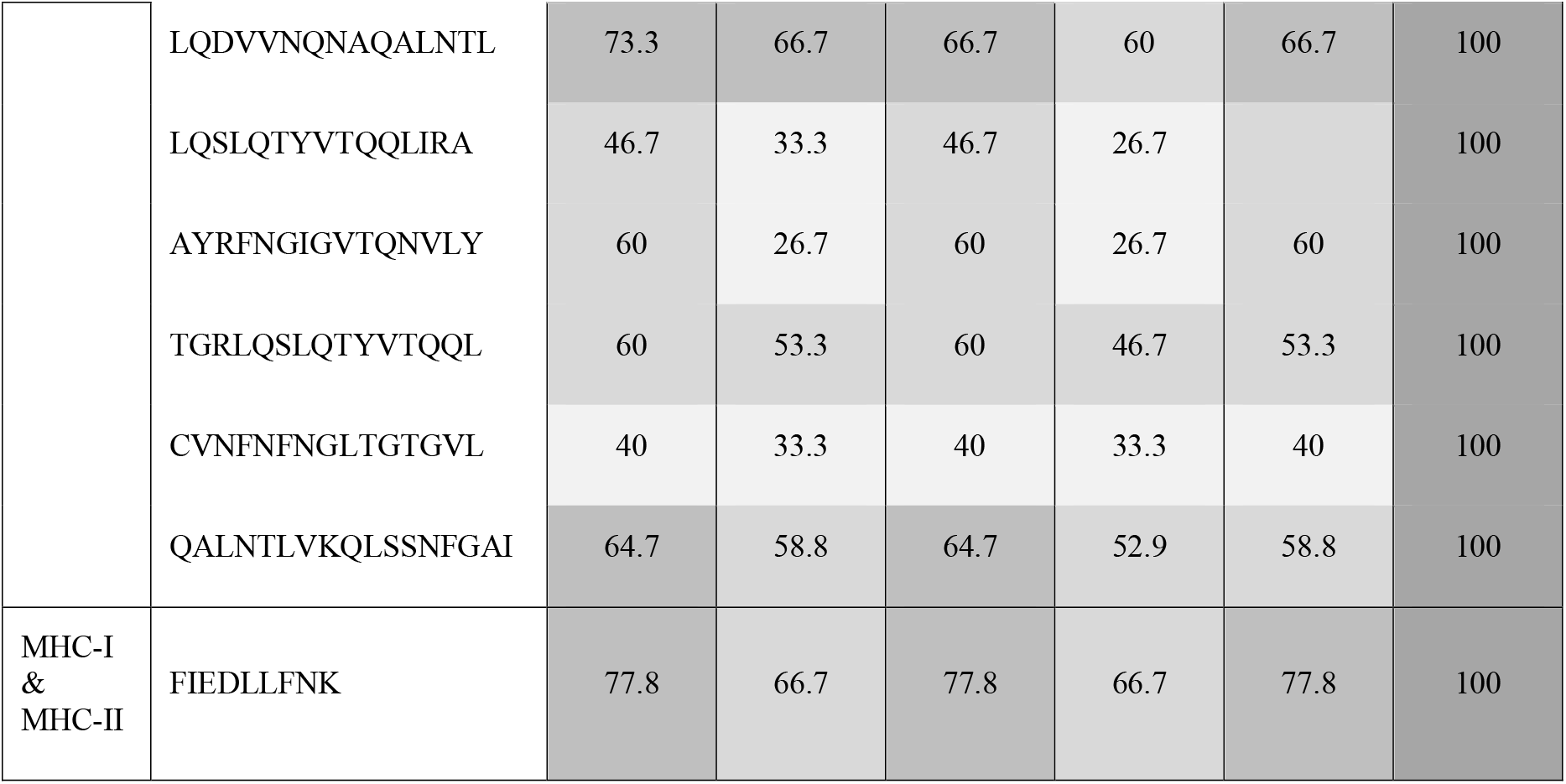
SARS-CoV-2 predicted T cell epitopes identity (%) shared with other coronaviruses.

## 4 Discussion

The emergence of SARS-CoV-2 as a human pathogen has drastically changed the traditional paradigm for vaccine development. For the first time in history hundreds of vaccine candidates have emerged and are currently being developed and tested at incredible speed (28) (https://www.covid-19vaccinetracker.org/). The urgent need for a vaccine means that development is being carried out without comprehensive knowledge of the virus pathogenesis, host response to infection, potentially detrimental immune responses and protective immunity. To keep up with vaccine demand around the world, more than one vaccine will be required and it’s highly possible that first generation vaccines will be far from perfect. In addition, we still lack a deep understanding of what is considered a protective immune response with most vaccine candidates focusing on neutralising antibodies, although the evidence suggests cellular immunity might play a critical role.

Through the literature review and epitope prediction exercises carried out in this project we were able to compile 82 potential linear B cell epitopes in the spike glycoprotein which could explain crossreactive responses reported in the literature. Twenty-seven novel epitopes were predicted, with the majority of them situated in the RBD and the NTD (Figure 2B). Recent studies have reported the identification of neutralising antibodies that target the NTD (29), therefore we consider the novel epitopes predicted in this study require to be further investigated as potential neutralising targets. Discontinuous B cell epitopes were also predicted in this study by using two different web-based tools. Even though ElliPro and DiscoTope yielded the prediction of epitopes in similar regions of the spike glycoprotein, there are limitations in the accuracy of these predictions given the nature of conformational epitopes. As more experimental data is generated, predictions of conformational epitopes for the SARS-CoV-2 spike glycoprotein will become more refined and precise. Despite the limitations, we believe that investigating potential neutralising antibodies against the predicted residues should be pursued. Given that coronaviruses have a latent pandemic potential we were interested to see if amongst the epitopes compiled in this manuscript there were conserved epitopes between the SARS-CoV-2 spike glycoprotein and other relevant human coronaviruses. We identified only 7 linear B cell epitopes that share certain identity between the SARS-CoV-2 spike glycoprotein and other coronaviruses. The percentages of identity between peptides ranged from 19 to 100%, with only one epitope with at least 60% identity between SARS-CoV-2 and all the other coronaviruses. Recent studies have shown that anti-spike antibodies generated in response to SARS-CoV infection recognise the spike glycoprotein of SARS-CoV-2 and vice versa, suggesting antigenic similarities between the spike glycoprotein of these two viruses. However, these cross reactive antibodies did not show any neutralising activity against other than the virus that caused the infection (30). Given that SARS-CoV and SARS-CoV-2 share the highest similarity it is highly unlikely that these antibodies could have neutralising activity against any of the other human coronaviruses. Another study showed that memory B cells from convalescent patients once infected with SARS-CoV, produce a repertoire of monoclonal antibodies that cross-neutralise SARS-CoV-2 while they showed no binding to OC43 or MERS spike glycoproteins (31). A different report showed that antibody responses against infection with seasonal human coronaviruses elicit neutralising antibodies against SARS-CoV-2, however responses were measured short after infection occurred and therefore there is no information about the potential longevity of such responses (32). All these observations show that cross-reactive humoral responses between coronaviruses are plausible, however it is still not clear whether they are robust enough to provide protection against infection and if they are long-lasting.

On the other hand a total of 66 potential T cell epitopes were predicted in this study, with 30 being MHC-I restricted, 32 MHC-II restricted, and 9 predicted to be promiscuous T cell epitopes. Even though most vaccines in current development are focusing on the generation of neutralising antibodies against the virus as a correlate of protection, recent studies strongly suggest a key role of T cell responses in controlling the infection (24, 33, 34). A study in mild and severe COVID-19 patients showed that in result to infection individuals developed SARS-CoV-2 specific T cells with central memory and effector memory phenotype, but more importantly mild cases generated higher frequencies of multi-cytokine producing CD8+ T cells (34). Strong memory T cell responses specific to SARS-CoV-2 have also been detected in samples from individuals who had mild or even asymptomatic infections, interestingly some in the absence of antibody responses (33). A study by Grifoni and collaborators reported T cells respond to stimulation with SARS-CoV-2 peptides in samples collected prior the current pandemic started, suggesting infections with seasonal coronaviruses elicit memory T cell responses that cross-react with SARS-CoV-2. Pending experimental confirmation, our work provides a panel of 38 T cell epitopes that might be responsible for these cross-reactive responses (Table 2). Together these observations strongly support the hypothesis that the role of cellular responses in controlling infection by SARS-CoV-2 might be as important as antibody responses, and could provide protection in the absence of humoral responses. In this context, the identification of potential protective T cell epitopes gains relevance in the design of next generation SARS-CoV-2 vaccines.

In conclusion, we were able to identify T cell epitopes from the SARS-CoV-2 glycoprotein that were found to have certain degree of identity between SARS-CoV-2 and the other human coronaviruses. The percentages of identity for these T cell epitopes were considerably higher to those found for B cell epitopes (Tables 1 & 2). Experimental evidence suggests robust T cell responses are generated in response to infection to SARS-CoV-2 and cross-reactive T cells circulate in the population prior SARS-CoV-2 exposure. The contribution of these responses towards protection remains to be further investigated in order to confirm our hypothesis. We hope that this manuscript will contribute to understand immune responses elicited in response to infection, and highlight the potential role of T cell epitopes in immunity and their influence in outbreak behaviour.

## 6 Author Contributions

CLM conceived the project, DLPO, SSRH, TRH and CLM designed the study; DLPO, SSRH, EAO and TRH acquired and analysed the data; DLPO, SSRH and TRH wrote the manuscript. EFO, LAP, ACV and CLM contributed in data interpretation. All authors read, revised and approved the submitted version of the manuscript.

## 8 Acknowledgments

We would like to thank Guillermo Ramón Torres for kindly facilitating the use of PyMOL^®^ software in order to build the 3D models presented in this manuscript.

## 12 Data Availability Statement

The datasets generated and analyzed for this study can be found in the bioRxv repository

