## Supplementary information for "Bioinformatic analysis of shared B and T cell epitopes amongst relevant coronaviruses to human health: Is there cross-protection?"

Supplementary Material

### 1 Supplementary Tables

| Supplementary Table 1. SARS-CoV-2 spike protein linear B cell epitopes | | |
| --- | --- | --- |
| **Sequence** | **Position** | **Source** |
| DVVNQNAQALNTLVKQL | 950-966 | (1) |
| **EIDRLNEVAKNLNESLIDLQELGKYEQY^*^**  EVAKNLNESLIDLQELG^*^ | 1182-1209 | (1) |
| FIEDLLFNKVTLADAGF | 817-833 | (1) |
| GAALQIPFAMQMAYRFN | 891-907 | (1) |
| GAGICASY | 667-674 | (1) |
| AISSVLNDILSRLDKVE | 972-988 | (1) |
| GSFCTQLN | 757-764 | (1) |
| ILSRLDKVEAEVQIDRL | 980-996 | (1) |
| AMQMAYRF | 899-906 | (1) |
| **KNHTSPDVDLGDISGIN^*^**  KNHTSPDVDLG | 1157-1173 | (1), this study |
| MAYRFNGIGVTQNVLYE | 902-918 | (1) |
| **RASANLAATKMSECVLGQSKRVD**  AATKMSECVLGQSKRVD  RASANLAATKMSECVLG | 1025-1041 | (1) |
| PFAMQMAYRFNGIGVTQ | 897-913 | (1) |
| QALNTLVKQLSSNFGAI | 957-973 | (1) |
| QLIRAAEIRASANLAAT | 1011-1027 | (1) |
| QQFGRD | 563-568 | (1) |
| **RLITGRLQSLQTYVTQQLIRAAEIR**  SLQTYVTQQLIRAAEIR  LQSLQTYVTQQLIRAAEI  RLITGRLQSLQTYVTQQ  VEAEVQIDRLITGRLQSL  TGRLQSLQTYVTQQL | 995-1019 | (1), (2) |
| CKFDEDDSEPVLKGVKLHYT | 1254-1273 | (1) |
| **TVYDPLQPELDSFKEEL**  VYDPLQPELDSF | 1133-1149 | (1), this study |
| **FRVQPTESVRFPNITNLCPFGEVFNATRFASVYAWNRKRISNCVA**^*^  FGEVFNAT^*^  VRFPNITNLCPFGEVFN^*^  FRVQPTESVRFPNITNLCPFGE^*^  FGEVFNATRFASVYA^*^  FPNITNLCPFGEVFNATRFASVYAWNRKRISNCVA^*^ | 327-343 | (1), (3), (2), this study |
| DDSEPVLKGVKLHYT | 1259-1273 | (1) |
| DLGDISGINASVVNIQK^*^ | 1165-1181 | (1) |
| **MFVFLVLLPLVSSQCVNLTTRTQLPPAYTNSFTRGVY**^*^  CVNLTTRTQLPPAYTN ^*^  VLLPLVSSQCVNLTTRTQLPPAYTN^*^  MFVFLVLLPLVSSQC  FVFLVLLPLVSSQCV  SQCVNLTTRTQLPPAYTNSFTRGVY^*^ | 14-30 | (4), (3), (2), this study |
| **FSNVTWFHAIHVSGTNGTKRFDN^*^**  NVTWFHAIHVSGTNGT^*^  VTWFHAIHVSGTNGTKRFDN^*^ | 60-76 | (4), (3), this study |
| **DPFLGVYYHKNNKSWME SEFRVYSSANNCTFEYVSQPFLM^*^**  YYHKNNKSWMESEFRVYSSANNCTFEYVSQPFLM^*^  DPFLGVYYHKNNKSWME^*^ | 142-175 | (3), this study |
| MDLEGKQGNFKNL | 175-187 | This study |
| **IYSKHTPINLVRDLPQGFS**  IYSKHTPIN  KHTPINLVRDLPQGFS | 204-214 | (3), this study |
| **RSYLTPGDSSSGWTAGAAAYLLKYNENGTIT^*^**  DSSSGWTAGAAAYYVG  TPGDSSSGWTA | 244-284 | (3), (2), this study |
| **DAVDCALDPLSETKCTLKSFTVEKGIYQTSN**  KGIYQTSN  KSFTVEKGIYQTSNFRVQP | 285-315 | (1, 3), this study |
| **YNSASFSTFKCYGVSPTKLNDLCFT**  SFSTFKCYGVSPTKLNDL | 367-391 | (4), this study |
| RSSVLHSTQD | 43-52 | (3) |
| VYFASTEKSNII | 89-99 | (3) |
| GTTLDSKTQSLLIVNNATNVVIKVC^*^ | 105-129 | (3) |
| IYSKHTPIN | 201-209 | (3) |
| DLPQGFSALEPLVDLPIGINITRFQTLLALH^*^ | 213-243 | (3) |
| **P YRVVVLSFELLHAPATVCG**  PYRVVVLSFELLHAP  YRVVVLSFELLHAPATVCG | 505-519  506-524 | (2) |
| YVGYLQPRTFLLKYN | 264-278 | (2) |
| IQDSLSSTASALGKL | 932-946 | (2) |
| **LFRKSNLKPFERDISTEIYQ**  VEGFNCYFPLQ | 453- 471  481-491 | (2) |
| GDEVRQIAPGQTGKIADYNYKLP | 404-426 | This study |
| ELLHAPATVCGPKKSTNLVKN | 514-535 | (2), this study |
| SNKKFLPF | 555-562 | This study |
| NCTEVPVAIHADQLTPT^*^ | 616-632 | This study |
| RVYSTGSNVFQ | 634-644 | This study |
| VNNSYECDIPI^*^ | 656-666 | This study |
| ASYQTQTNSPRRARSVASQ | 672-690 | This study |
| **IIAYTMSLGAENSVAYSNN^*^**  YTMSLGAENSVAYSNN^*^  IIAYTMSLGAENSVA | 690-708 | (2), this study |
| EQDKNTQ | 773-779 | This study |
| KQIYKTPPIKDFGGF | 786-800 | This study |
| PDPSKPSK | 807-814 | This study |
| LADAGFIKQYGDCLG | 828-842 | This study |
| GQSKRVDFC | 1035-1043 | This study |
| RNFYEPQIITTD | 1107-1118 | This study |
| SCCKFDEDDSEPVLKG | 1252-1267 | This study |
| MDLEGKQGNFKNL | 177-189 | This study |
| NGTITD^*^ | 282-287 | This study |
| VRQIAPGQTGKIAD | 407-420 | This study |
| YKLPDD | 423-428 | This study |
| NNLDSKVGG | 439-447 | This study |
| YQAGSTPCNGV | 473-483 | This study |
| YGFQPTNGVGYQ | 495-506 | This study |
| RDIADTTDAVRDPQ | 567-580 | This study |
| VITPGTNTSN | 597-606 | This study |
| QTQTNSPRRARSV | 675-687 | This study |
| VEQDKNTQE | 772-780 | This study |
| IYKTPPIKDF | 788-797 | This study |
| ILPDPSKPSKRS | 805-816 | This study |
| **GYHLMSFPQSAPHGVVFLHVTYVPAQEKNFTT**  GVVFLHVTYVPAQEK  PAQEKNFTT^*^ | 1056-1074 | (2), this study |
| **DVVIGIVNNTVYDPLQPELDSF^*^**  DVVIGIVNNTVYDPLQPE^*^  VYDPLQPELDSF | 1124-1145 | (2), this study |
| ^*^ Glycopeptide  Peptides in bold letters are those which contain all the overlapping epitopes predicted by various studies | | |

| Supplementary Table 2. SARS-CoV-2 spike protein T cell epitopes | | | | |
| --- | --- | --- | --- | --- |
| **Sequence** | **Position** | **Source** | **HLA affinity** | **MHC binding** |
| RGWIFGTTLDSKTQSLL | 100-116 | (5) | HLA-DRB1*04:01 | II |
| CTFEYVSQPFLMD  YVSQPFLMD | 164-176 | (5), this study | HLA-DRB1*04:01, HLA-DPA1*02:02/DPB1*02:02, HLA-DPA1*02:01/DPB1*02:01, HLA-DPA1*03:01/DPB1*23:01 | II |
| QPFLMDLEGKQGN | 171-183 | (5) | HLA-DRB1*04:01 | II |
| TRFQTLLALHRSYLTPGDSSSGW  FQTLLALHR  LALHRSYLT | 234-256 | (5), this study | HLA-DRB1*04:01,  HLA-DRB1*01:01,  HLA-DPA1*01:03/DPB1*03:01,  HLA-DPA1*02:02/DPB1*02:02,  HLA-DPA1*03:01/DPB1*23:01,  HLA-DPA1*02:01/DPB1*02:01 | II |
| KSFTVEKGIYQTSNFRVQ  FTVEKGIYQ | 302-319 | (5), this study | HLA-DRB1*04:01, HLA-DRB1*07:01 | II |
| SASFSTFKCYGVSPTKL | 302-319 | (5) | HLA-DRB1*0701, HLA-DR8 | II |
| KLPDDFTGCV | 422-431 | (5) | HLA-A*02:01 | I |
| NLDSKVGGNYNYLYRLFR  NYNYLYRLF | 438-456 | (5), this study | HLA-A*24:02  HLA-A*23:01 | I |
| YLYRLFRKSNLKPFERDI | 449-466 | (5) | HLA-DRB1*04:01 | II |
| KPFERDISTEIYQ | 461-468 | (5) | HLA-DRB1*04:01 | II |
| QSIIAYTMSLGAENSVAY  SIIAYTMSL  LGAENSVAY | 688-705 | (5), this study | HLA-DRB1*04:01, HLA-DRB*07:01, HLA-A*02:01, HLA-B*35:01 | II |
| TECSNLLLQYGSFCTQL | 745-761 | (5) | HLA-DR8 | II |
| VKQIYKTPPIKDFGGFNF^*^ | 783-800 | (5) | HLA-DRB1*04:01 | II |
| DSLSSTASALGKLQDVV | 934-950 | (5) | HLA-DRB1*04:01 | II |
| ALNTLVKQL | 956-964 | (1, 5) | HLA-A*02:01 | I |
| VLNDILSRL  SVLNDILSR | 974-983 | (1, 5), this study | HLA-A*02:01  HLA-A*11:01, HLA-A*31:01, HLA-A*68:01 | I |
| LITGRLQSL | 994-1001 | (1, 5) | HLA-A2, HLA-A*02:01 | I |
| QLIRAAEIRASANLAATK  IRASANLAA | 1008-1025 | (1, 5), this study | HLA-DRB1*04:01,  HLA-DRB1*01:01,  HLA-DPA1*01:03/DPB1*03:01,  HLA-DQA1*05:01/DQB1*03:01 | II |
| HWFVTQRNFYEPQII | 1098-1112 | (5) | HLA-DRB1*04:01 | II |
| RLNEVAKNL | 1182-1190 | (5) | HLA*02:01 | I |
| NLNESLIDL^*^ | 1189-1197 | (5) | HLA*02:01 | I |
| EPVLKGVKL | 949-957 | (4) | HLA-B*07:02 | I |
| VVNQNAQAL | 949-957 | (4) | HLA-B*07:02 | I |
| YLQPRTFLL | 267-275 | (4), this study | HLA-A*02:01, HLA-A*02:06, HLA-B*08:01,  HLA-DRB1*01:01,  HLA-DPA1*02:02/DPB1*02:02,  HLA-DPA1*02:01/DPB1*02:01,  HLA-DPA1*03:01/DPB1*23:01 | I, II |
| FQFCNDPFL | 133-141 | This study | HLA-A*02:06, HLA-A*02:01 | I |
| QYIKWPWYI | 1208-1216 | This study | HLA-A*23:01 | I |
| LPFNDGVYF | 83-92 | This study | HLA-B*35:01 | I |
| LTDEMIAQY | 864-873 | This study | HLA-A*01:01 | I |
| FIAGLIAIV | 1220-1229 | (1), this study | HLA-A*02:01, HLA-A*02:02, HLA-A*02:03, HLA-A*02:06, HLA-A*68:02, HLA-A2 | I |
| QYIKWPWYI | 1207-1216 | This study | HLA-A*24:02 | I |
| SPRRARSVA | 680-688 | This study | HLA-B*07:02 | I |
| FTISVTTEI | 717-726 | This study | HLA-A*02:06, HLA-DRB1*07:01, HLA-DPA1*03:01/DPB1*23:01  HLA-DPA1*02:02/DPB1*02:02  HLA-DPA1*03:01/DPB1*23:01 | I, II |
| GVYFASTEK | 88-95 | This study | HLA-A*11:01,  HLA-A*03:01 | I |
| VVFLHVTYV | 1060-1068 | (1), this study | HLA-A*02:01, HLA-A*02:02, HLA-A*02:03, HLA-A*02:06, HLA-A*68:02 | I |
| ASANLAATK | 1020-1029 | (1), this study | HLA-A*11:01, HLA-A*03:01, HLA-A*68:01, HLA-A*31:01 | I |
| YEQYIKWPW | 1206-1215 | (1), this study | HLA-B*44:02, HLA-B*18:01, HLA-B*44:03, HLA-B*40:01, HLA-B*45:01 | I |
| SANNCTFEY^*^ | 162-170 | This study | HLA-B*35:01 | I |
| AEIRASANLA | 1016-1025 | (1), this study | HLA-B*45:01, HLA-B*40:01, HLA-B*44:02, HLA-B*44:03, HLA-B*40:02 | I |
| NATRFASVY^*^ | 343-351 | This study | HLA-B*35:01 | I |
| YQDVNCTEV^*^ | 611-620 | This study | HLA-A*02:06 | I |
| VTYVPAQEK | 1065-1074 | This study | HLA-A*11:01, HLA-A*03:01 | I |
| SVTTEILPV | 721-729 | This study | HLA-A*02:06 | I |
| TLKSFTVEK | 302-311 | This study | HLA-A*11:01, HLA-A*03:01 | I |
| LPFFSNVTW^*^ | 56-65 | This study | HLA-B*35:01 | I |
| GTHWFVTQR | 1099-1107 | This study | HLA-A*11:01 | I |
| WTAGAAAYY  GWTAGAAAY | 256-264 | (4), this study | HLA-A*01:01, HLA-B*35:01, HLA-B*07:02, HLA-DQA1*05:01/DQB1*03:01 | I, II |
| FLHVTYVPA | 1062-1071 | This study | HLA-A*02:01, HLA-A*02:06 | I |
| TLADAGFIK | 827-835 | This study | HLA-A*11:01 | I |
| ASVYAWNRK | 348-357 | This study | HLA-A*11:01 | I |
| VTWFHAIHV | 62-71 | This study | HLA-A*02:06 | I |
| RLDKVEAEV | 982-991 | (1), this study | HLA-A*02:01, HLA-A*02:02, HLA-A*02:06, HLA-A*02:03, HLA-A*68:02 | I |
| WPWYIWLGF | 1212-1221 | This study | HLA-B*35:01 | I |
| FTISVTTEI | 717-726 | This study | HLA-A*02:01 | I |
| QPTESIVRF | 320-329 | This study | HLA-B*35:01 | I |
| DSFKEELDKY | 1146-1155 | (1) | HLA-A*26:01, HLA-A*29:02, HLA-A*30:02, HLA-A*01:01 | I |
| EDLLFNKVTLADAGF | 819-833 | (1) | HLA-DRB1*01:01 | II |
| AEVQIDRLI | 989-997 | (1) | HLA-B*40:02, HLA-B*44:02, HLA-B*44:03, HLA-B*45:01 | I |
| AEVQIDRLIT | 989-998 | (1) | HLA-B*40:02, HLA-B*44:03, HLA-B*45:01, HLA-B*40:01, HLA-B*44:02 | I |
| FPNITNLCPF^*^ | 329-338 | (1) | HLA-B*35:01, HLA-B*51:01, HLA-B*53:01, HLA-B*07:02, HLA-B*54:01 | I |
| GAALQIPFAMQMAYR | 891-905 | (1) | HLA-DRB1*01:01 | II |
| GLIAIVMVTI | 1223-1232 | (1) | HLA-A*02:02, HLA-A*02:03, HLA-A*02:01, HLA-A*02:06, HLA-A*68:02 | I |
| GRLQSLQTY | 999-1007 | (1) | HLA-B*27:05 | I |
| GSFCTQLNR | 757-765 | (1) | HLA-A*11:01, HLA-A*31:01, HLA-A*68:01, HLA-A*03:01 | I |
| GVVFLHVTY | 1059-1067 | (1) | HLA-A*11:01 | I |
| GWTFGAGAALQIPFA  WTFGAGAAL | 885-899 | (1), this study | HLA-DRB1*01:01,  HLA-DQA1*05:01/DQB1*03:01 | II |
| GYQPYRVVVL | 504-513 | (1) | HLA-A*23:01, HLA-A*24:02, HLA-A*29:02 | I |
| IDRLITGRLQSLQTY | 993-1007 | (1) | HLA-DRB1*01:01 | II |
| IEDLLFNKVTLADAG | 818-832 | (1) | HLA-DRB1*01:01 | II |
| IGAGICASY | 666-674 | (1) | HLA-A*29:02, HLA-A*30:02 | I |
| IITTDNTFV | 1114-1122 | (1) | HLA-A*02:01 | I |
| ISGINASVVNIQKEI^*^ | 1169-1183 | (1) | HLA-DRB1*01:01 | II |
| LDKYFKNHTSPDVDL^*^ | 1152-1166 | (1) | HLA-DRB1*01:01 | II |
| APHGVVFLHV | 1056-1065 | (1) | HLA-B*07:02, HLA-B*54:01, HLA-B*35:01, HLA-B*53:01 | I |
| LGDISGINASVVNIQ^*^  ISGINASVV^*^ | 1166-1180 | (1), this study | HLA-DRB1*01:01,  HLA-DQA1*05:01/DQB1*03:01 | II |
| LGFIAGLIAIVMVTI  WLGFIAGLIAIVMVT  IAGLIAIVM  WLGFIAGLI  LGFIAGLIA | 1217-1231 | (1), this study | HLA-DRB1*01:01,  HLA-DPA1*02:01/DPB1*02:01,  HLA-DQA1*05:01/DQB1*03:01,  HLA-DPA1*03:01/DPB1*23:01 | II |
| LIDLQELGKY | 1197-1206 | (1) | HLA-A*30:02, HLA-A*01:01, HLA-A*26:01, HLA-A*29:02 | I |
| LLFNKVTLA | 821-829 | (1) | HLA-A*02:01, HLA-A*02:02, HLA-A*02:03, HLA-A*02:06, HLA-A*68:02 | I |
| LLLQYGSFC  LLQYGSFCT | 752-760 | (1) | HLA-A*02:01 | I |
| LNTLVKQLSSNFGAI  VKQLSSNFG | 959-973 | (1), this study | HLA-DRB1*01:01,  HLA-DQA1*05:01/DQB1*03:01 | II |
| LQDVVNQNAQALNTL | 948-962 | (1) | HLA-DRB1*01:01 | II |
| LQIPFAMQM | 894-902 | (1) | HLA-B*15:01, HLA-C*15:02, HLA-B*40:01, HLA-B*58:01 | I |
| LQSLQTYVTQQLIRA  LQTYVTQQLIRAAEI  QQLIRAAEIRASANL  QTYVTQQLIRAAEIR  LQSLQTYVT | 1001-1033 | (1), this study | HLA-DRB1*01:01,  HLA-DPA1*02:02/DPB1*02:02,  HLA-DPA1*03:01/DPB1*23:01,  HLA-DPA1*02:01/DPB1*02:01 | II |
| AQALNTLVK | 956-964 | (1) | HLA-A*11:01, HLA-A*03:01, HLA-A*31:01, HLA-A*68:01 | I |
| AQKFNGLTVLPPLLT  FNGLTVLPP | 852-866 | (1), this study | HLA-DRB1*01:01,  HLA-DPA1*01:03/DPB1*03:01,  HLA-DPA1*03:01/DPB1*23:01,  HLA-DPA1*02:02/DPB1*02:02,  HLA-DPA1*02:01/DPB1*02:01 | II |
| MTSCCSCLK | 1237-1245 | (1) | HLA-A*11:01, HLA-A*31:01, HLA-A*33:01, HLA-A*68:01, HLA-A*03:01 | I |
| NLNESLIDL^*^ | 1192-1200 | (1) | HLA-A*02:01 | I |
| PCSFGGVSVITPGTN^*^ | 589-603 | (1) | HLA-DRB1*01:01 | II |
| PYRVVVLSF | 507-515 | (1) | HLA-A*23:01, HLA-A*24:02, HLA-A*01:01, HLA-A*26:01 | I |
| QELGKYEQYI | 1201-1210 | (1) | HLA-B*44:02, HLA-B*44:03, HLA-B*40:02 | I |
| QPYRVVVLSF | 506-515 | (1) | HLA-B*07:02, HLA-B*53:01 | I |
| RLNEVAKNL | 1185-1193 | (1) | HLA-A*02:01 | I |
| RLQSLQTYV | 1000-1008 | (1) | HLA-A*02:01, HLA-A*02:02, HLA-A*02:03, HLA-A*02:06, HLA-A*68:02 | I |
| RVDFCGKGY | 1039-1047 | (1) | HLA-A*30:02, HLA-A*01:01, HLA-A*03:01, HLA-B*15:01, HLA-B*27:05 | I |
| SEPVLKGVKL | 1261-1270 | (1) | HLA-B*40:01, HLA-B*40:02, HLA-B*44:02, HLA-B*44:03 | I |
| SFIEDLLFNK  FIEDLLFNK | 816-825 | (1), this study | HLA-A*11:01, HLA-A*31:01, HLA-A*33:01, HLA-A*68:01,  HLA-DPA1*02:02/DPB1*02:02,  HLA-DPA1*02:01/DPB1*02:01,  HLA-DPA1*03:01/DPB1*23:01 | I, II |
| AYRFNGIGVTQNVLY | 903-917 | (1) | HLA-DRB1*01:01 | II |
| SLIDLQELGK | 1196-1205 | (1) | HLA-A*03:01, HLA-A*11:01, HLA-A*31:01, HLA-A*68:01, HLA-A*33:01 | I |
| SSNFGAISSVLNDIL  FGAISSVLN | 967-981 | (1), this study | HLA-DRB1*01:01,  HLA-DPA1*02:02/DPB1*02:02,  HLA-DPA1*02:01/DPB1*02:01,  HLA-DPA1*03:01/DPB1*23:01 | II |
| TGRLQSLQTYVTQQL | 998-1012 | (1) | HLA-DRB1*01:01 | II |
| TQNVLYENQK | 912-921 | (1) | HLA-A*11:01,HLA-A*68:01 | I |
| CMTSCCSCLK | 1236-1245 | (1) | HLA-A*68:01, HLA-A*03:01, HLA-A*11:01, HLA-A*31:01, HLA-A*33:01 | I |
| VQIDRLITGR | 991-1000 | (1) | HLA-A*31:01, HLA-A*03:01, HLA-A*11:01, HLA-A*33:01, HLA-A*68:01 | I |
| VRFPNITNL^*^ | 327-365 | (1) | HLA-C*14:02, HLA-B*27:05 | I |
| VYDPLQPEL | 1137-1145 | (1) | HLA-A*24:02, HLA-A*29:02, HLA-A*30:02 | I |
| CVADYSVLY | 361-369 | (1) | HLA-A*01:01, HLA-A*26:01, HLA-A*29:02, HLA-A*30:02 | I |
| CVNFNFNGLTGTGVL  FNGLTGTGV | 538-552 | (1), this study | HLA-DRB1*01:01,  HLA-DQA1*05:01/DQB1*03:01 | II |
| DDSEPVLKGVKLHYT | 1259-1273 | (1) | HLA-DRB1*01:01 | II |
| DEDDSEPVL | 1257-1265 | (1) | HLA-B*40:01, HLA-B*40:02 | I |
| DKYFKNHTSPDVDLG^*^ | 1153-1167 | (1) | HLA-DRB1*01:01 | II |
| GAALQIPFAMQMAYRF | 891-906 | (1) | HLA-DRA*01:01, HLA- DRB1*07:01 | II |
| MAYRFNGIGVTQNVLY | 902-917 | (1) | HLA-DRB1*04:01 | II |
| QALNTLVKQLSSNFGAI | 957-973 | (1) | HLA-DRB1*04:01 | II |
| FLVLLPLVS | 4-12 | This study | HLA-DRB1*01:01 | II |
| VLSFELLHA | 512-521 | This study | HLA-DRB1*01:01 | II |
| FTISVTTEI | 717-726 | This study | HLA-DRB1*07:01 | II |
| KCVNFNFNG | 537-546 | This study | HLA-DRB1*01:01,  HLA-DQA1*05:01/DQB1*03:01 | II |
| ITRFQTLLA | 235-243 | This study | HLA-DRB1*01:01,  HLA-DPA1*02:02/DPB1*02:02  HLA-DPA1*02:01/DPB1*02:01  HLA-DPA1*03:01/DPB1*23:01  HLA-DPA1*01:03/DPB1*03:01 | II |
| YSVLYNSAS | 365-373 | This study | HLA-DRB1*01:01,  HLA-DPA1*02:01/DPB1*02:01,  HLA-DPA1*02:02/DPB1*02:02,  HLA-DPA1*03:01/DPB1*23:01,  HLA-DQA1*05:01/DQB1*03:01 | II |
| FTVEKGIYQ | 306-314 | This study | HLA-DRB1*07:01 | II |
| QIPFAMQMAYRFNGI  FAMQMAYRF | 898-907 | (1), this study | HLA-DRB1*01:01,  HLA-DPA1*01:03/DPB1*03:01,  HLA-DPA1*02:02/DPB1*02:02,  HLA-DPA1*03:01/DPB1*23:01,  HLA-DPA1*02:01/DPB1*02:01 | II |
| FCTQLNRAL | 759-767 | This study | HLA-DRB1*01:01,  HLA-DQA1*05:01/DQB1*03:01 | II |
| IAQYTSALL | 869-878 | This study | HLA-DRB1*01:01,  HLA-DQA1*05:01/DQB1*03:01 | II |
| FKNIDGYFK | 194-203 | This study | HLA-DRB1*01:01,  HLA-DPA1*02:02/DPB1*02:02,  HLA-DPA1*02:01/DPB1*02:01 | II |
| YFKIYSKHT | 200-208 | This study | HLA-DRB1*01:01 | II |
| LIVNNATNV^*^ | 117-126 | This study | HLA-DRB1*01:01,  HLA-DQA1*05:01/DQB1*03:01 | II |
| FNATRFASV^*^ | 342-351 | This study | HLA-DPA1*02:02/DPB1*02:02,  HLA-DPA1*02:01/DPB1*02:01,  HLA-DPA1*03:01/DPB1*23:01,  HLA-DQA1*05:01/DQB1*03:01 | II |
| FSNVTWFHA^*^ | 59-68 | This study | HLA-DPA1*02:02/DPB1*02:02,  HLA-DQA1*05:01/DQB1*03:01 | II |
| RVVVLSFEL | 509-517 | This study | HLA-DPA1*02:02/DPB1*02:02 | II |
| PIGAGICAS | 665-674 | This study | HLA-DQA1*05:01/DQB1*03:01 | II |
| SSGWTAGAA | 254-263 | This study | HLA-DQA1*05:01/DQB1*03:01 | II |
| LLAGTITSG | 877-885 | This study | HLA-DQA1*05:01/DQB1*03:01 | II |
| ^*^ Glycopeptide | | | | |

### 2 Supplementary Figures

**Supplementary Figure 1.** Linear B cell epitopes of the SARS-CoV-2 spike glycoprotein displayed in the monomer representation side. Epitopes reported in the literature and assignment of functional domains (A), epitopes predicted with IEDB tools (B) and epitopes that are shared with other coronaviruses predicted or reported (C).

**Supplementary Figure 2.** T cell epitoprs of the SARS-CoV-2 spike glycoprotein displayed in the monomer representation side. Epitopes reported in the literature and assignment of functional domains (A), epitopes predicted with IEDB tools (B) and epitopes that are shared with other coronaviruses predicted or reported (C).
